# Longitudinal analysis reveals myeloid cell contributions to neuroPASC pathogenesis

**DOI:** 10.64898/2026.05.04.722452

**Authors:** Lu Tan, Shea Lowery, Abhishek K. Verma, Andrew Thurman, Alan Sariol, Cori Fain, John Harty, Stanley Perlman

**Affiliations:** Department of Microbiology and Immunology, University of Iowa, Iowa City, IA 52242, USA; Department of Internal Medicine, University of Iowa, Iowa City, IA 52242, USA; Department of Medicine, Washington University School of Medicine, St. Louis, MO 63110, USA; Department of Pathology, University of Iowa, Iowa City, IA 52242, USA

## Abstract

Neurological and neuropsychiatric symptoms, collectively termed neuroPASC, are among the most prevalent Post-Acute Sequelae of COVID-19 (PASC). Neuroinflammation – particularly microglia reactivity – has been implicated in neuroPASC. Current insights are largely derived from post-mortem tissues of acutely infected patients and may not reflect PASC-related neuropathology. We previously established a PASC model in which SARS-CoV-2-infected mice developed persistent behavioral alterations and prolonged neuroinflammation for up to 120 days post-infection (dpi) in the absence of viral neuroinvasion. Here, we extended these results to a longitudinal single-cell RNA sequencing analysis of brain immune cells collected at 0, 6, 30, and 100 dpi. We identified a coordinated contribution of infiltrating and resident myeloid cells to the initiation and persistence of neuroinflammation. In specific, microglia displayed sustained expansion of subclusters characterized by inflammatory, stress response, and metabolic signatures. Border-associated macrophages upregulated monocyte attractants during acute infection. Concurrently, monocytes and neutrophils showed marked brain recruitment and mounted transient inflammatory responses at 6 dpi, potentially triggering long-term microglial reactivity. Together, these findings provide a high-resolution atlas of brain myeloid immune dynamics during neuroPASC and highlight a central role for microglia in sustaining chronic neuroinflammation.

## INTRODUCTION

COVID-19, caused by the severe acute respiratory syndrome coronavirus 2 (SARS-CoV-2), led to substantial morbidity from 2020-2023 and still causes 20,000-50,000 deaths in the USA per year, mostly in aged and immunocompromised people^1^. While acute disease has been ameliorated in most populations, it has become clear that many previously infected individuals develop long term effects of the infection (post-acute sequelae of COVID-19, PASC), which encompass a range of multisystem disorders. Owing to its complexity and prevalence, PASC poses substantial impacts on individual lives along with broad economic and healthcare consequences^2^. Neurological and psychiatric manifestations – collectively termed neuroPASC – are among the most common sequelae, affecting both previously hospitalized and non-hospitalized patients and persisting for at least three years in some cases^3^. Typical symptoms/signs include, but are not limited to, brain fog, fatigue, dizziness, olfactory dysfunction, memory problems, abnormal movements, epilepsy/seizure, ischemic stroke, and mixed anxiety disorder^3^. Accumulating evidence implicates multifaceted pathological processes in these COVID-19-associated neurological dysfunctions. Neuroimaging studies have revealed reductions in grey matter volume in patients with acute and prolonged cognitive declines^4, 5^, accompanied by sustained elevations of brain injury markers in serum^5, 6^. Cerebral microvascular obstruction^7^, blood-brain barrier (BBB) disruption^8, 9^, and neuroinflammation^8, 10, 11, 12^ have been documented in patient cohorts. Histological, proteomic, and transcriptomic analyses of post-mortem brains further demonstrate broad cellular perturbations in the central nervous system (CNS), involving neurons, microglia, oligodendrocytes, astrocytes, and T cells, among others^10, 12, 13, 14, 15^. Notably, although SARS-CoV-2 infection has been reported in human brain organoids^16, 17, 18^ and pluripotent stem cell-derived neural cells^16, 19^, the detection of viral RNA or protein within the human CNS has been reported in only a minority of studies^10, 12, 18, 20, 21, 22^.

Among various observed pathological alterations, neuroinflammation appears to play a central role. It has been linked to or promoted by systemic inflammation^9, 12^, fibrin deposition within the brain parenchyma^23^, and possibly, persistence of SARS-CoV-2 spike protein along the skull-meninges-brain axis^24^. Microglia, yolk sac-derived, CNS parenchymal resident myeloid cells, are key mediators of neuroinflammation across diverse contexts such as neurodegenerative diseases and aging^25, 26, 27^. Their reactivity has been consistently reported in COVID-19 patients and animal models, as evidenced by morphological changes, upregulation of activation markers, and increased cell numbers^8, 10, 12, 23, 28, 29^. Their underlying transcriptional reprogramming has also been profiled using single-cell and single-nucleus RNA sequencing (scRNA-seq and snRNA-seq)^12, 13, 14, 15, 30, 31^. However, most existing analyses were performed on post-mortem patient samples with heterogeneous disease courses and comorbidities^12, 13, 14, 15, 30^ and are often restricted to specific brain regions – primarily the cortex or brainstem^12, 13, 14, 15, 30, 31^. Given the broad spectrum of COVID-19-associated neurological symptoms, it is likely that additional brain regions are also affected but remain insufficiently characterized.

Border-associated macrophages (BAMs) constitute another population of brain-resident sentinel cells localized to meninges, choroid plexus (CP), and perivascular spaces^32^. Alongside hematogenous monocyte-derived macrophages entering the CNS under pathological conditions^32^, they exhibit both redundant and non-redundant functions relative to microglia. For example, in autoimmune demyelinating diseases such as multiple sclerosis (MS), both microglia and macrophages engulf myelin debris, whereas microglia additionally promote oligodendrocyte differentiation and remyelination^33, 34, 35^. However, possible contributions of BAMs and monocyte-derived macrophages during SARS-CoV-2 infection remain undefined. Of note, outside the CNS, monocytes and macrophages are potent drivers of systemic inflammation and COVID-19 pathogenesis^36, 37, 38^. Similarly, neutrophils have been implicated in acute COVID-19 severity and post-COVID-19 pulmonary sequelae^39, 40, 41^, but it is unclear whether these responses extend to the CNS.

Altogether, a comprehensive and longitudinal characterization of whole-brain myeloid cell dynamics following peripheral SARS-CoV-2 infection is still lacking. Given the scarcity of post-mortem samples, especially from neuroPASC patients, relevant animal models are essential. Because ancestral strains of SARS-CoV-2 cannot infect mice, we and others generated mouse-adapted viruses. We introduced a N501Y mutation into the spike protein using reverse genetics, allowing the virus to use mouse ACE2 for cellular entry^42, 43^. We then passaged the virus 30 times through mouse lungs to isolate a virulent, mouse-adapted SARS-CoV-2 (SARS2-N501Y_MA30_)^43^. Using this virus, we showed that mice develop anosmia during acute infection and persistent neuroinflammation with behavioural changes^44, 45^.

In this study, we used this experimental infection model to longitudinally assess the CNS inflammatory milieu of neuroPASC, up to 100 days post-infection (dpi). We found that inflammatory myeloid cell migration into the brain and antiviral responses were maximal at early times after infection and slowly decreased over the next few weeks. In contrast, microglial reactivity was detected at all time points post-infection. Importantly, these changes occurred independent of viral neuroinvasion. Together, our data provide a high-resolution atlas of brain immune responses during neuroPASC progression and underscore the predominant role of microglia in sustaining long-term neuroinflammation.

## RESULTS

### A neuroPASC mouse model

A sublethal dose (1000 PFU per mouse) of SARS2-N501Y_MA30_ was intranasally administered to 4-6-month-old C57BL/6 mice to establish a neuroPASC model, based on our previous results^45^. This age range was chosen because young C57BL/6 mice (6-10 weeks) are only modestly susceptible to SARS2-N501Y_MA30_ infection^43^, while mice greater than 4 months uniformly develop dosage-dependent clinical disease. Here, we showed that infected mice exhibited over 20% body weight loss by 5-7 dpi, followed by gradual recovery, although an approximate 10% weight loss persisted at 14 dpi (Fig. 1a). Meanwhile, more than one-quarter of animals succumbed to infection between 5 and 10 dpi (Fig. 1b). These data suggested that this model recapitulated more severe symptoms and a prolonged disease course compared with other reported long COVID mouse models^28, 29, 31^. As expected for a respiratory tract infection, high levels of viral RNA (vRNA) were detected in lungs at 2 dpi and remained detectable even at 60 dpi (Fig. 1c), although infectious viruses were no longer recoverable by 10 dpi. In contrast, vRNA was only transiently detected in the brain, declining to the mock level thereafter (Fig. 1c). This is consistent with our previous report demonstrating that a small number of infectious virus particles can be found in some mouse brains at 2 dpi^43^, likely as a consequence of viremia, since no virus-infected cells were observed in the brain^46^. Therefore, we consider productive CNS infection in our model to be unlikely.

**Figure 1.**
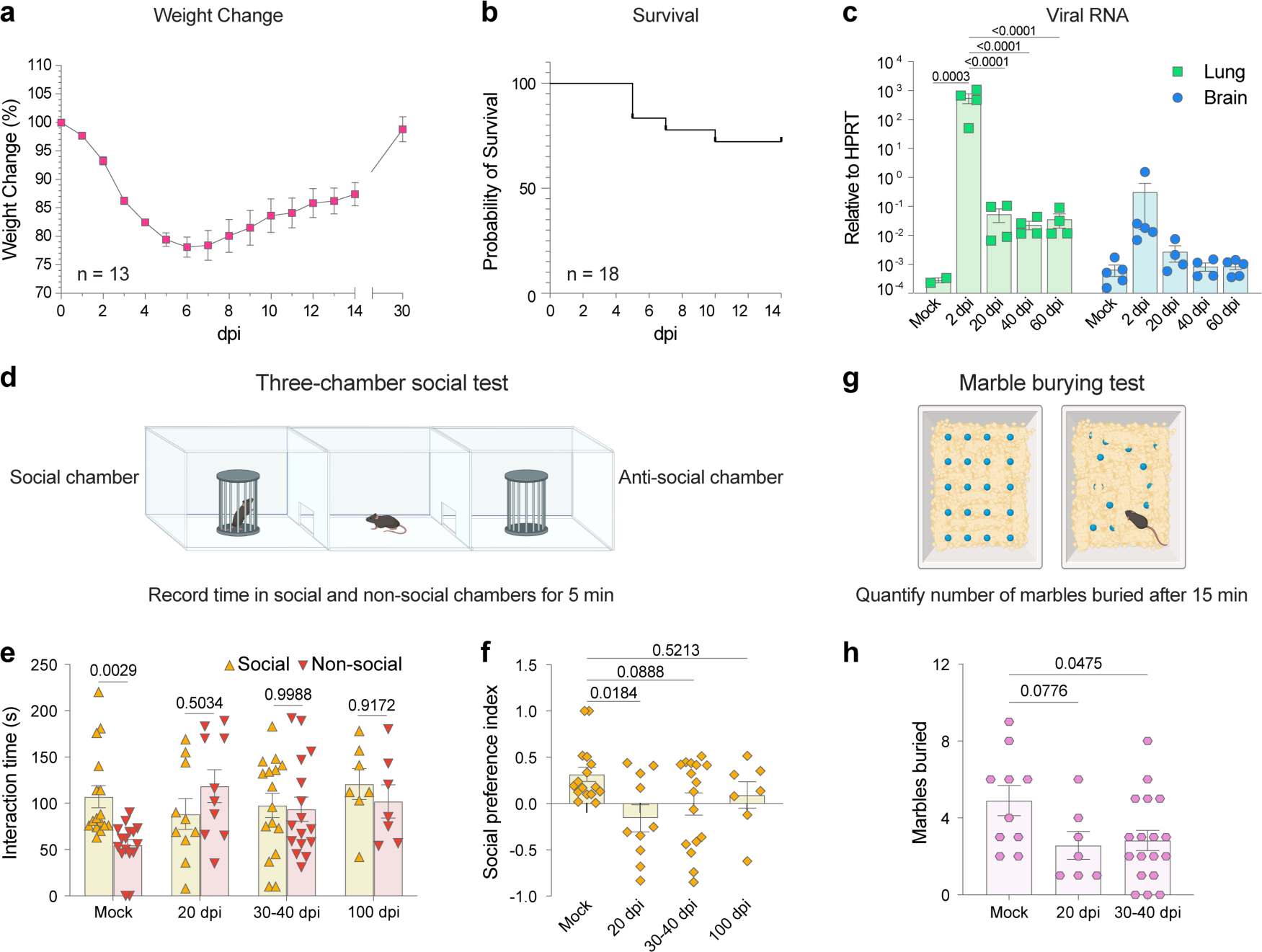
Long-term behavioural changes in a neuroPASC mouse model. **a** | Body weight change of C57BL/6 mice (4–6 months old) following intranasal infection with SARS2-N501Y_MA30_ (1000 PFU, n=13). **b** | Survival of infected mice (n=18). **c** | Viral RNA loads in lungs and brains at indicated post-infection time points. **d** | Schematic of the three-chamber social preference test. **e** | Time spent by mice in social (T_S_) versus non-social (T_NS_) chambers (mock: n = 16; 20 dpi: n = 10; 30–40 dpi: n = 16; 100 dpi: n = 7). **f** | Social preference index (I_SP_) of individual time points. I_SP_ was calculated as: I_SP_ = (T_S_ − T_NS_)/(T_S_ + T_NS_). **g** | Schematic of the marble burying test. **h** | Number of marbles buried (mock: n = 10; 20 dpi: n = 7; 30–40 dpi: n = 18). A marble was scored as buried if ≥ 60% of its surface was covered with bedding. All data are presented as mean ± standard error of the mean (SEM). Statistical significance in (**c**, **e**) was determined using ordinary two-way ANOVA with Sidak’s multiple comparisons test, and significance in (**f**, **h**) was assessed by ordinary one-way ANOVA with Dunnett’s multiple comparisons test. Non-significant *p* values in (**c**) are not shown.

Given the wide range of COVID-19-related CNS symptoms and signs^3^, we performed several behavioral tests to evaluate the clinical manifestations of neuroPASC. We previously reported motor dysfunction in SARS2-N501Y_MA30_-infected mice, as demonstrated by deficits in rotarod and open field tests^45^. Next, we assessed potential social impairment by conducting three-chamber social testing (Fig. 1d)^47^. Healthy (mock-infected) mice are considered social animals, and as expected, they spent more time exploring another mouse (“social stimulus”) than an empty chamber (“non-social stimulus”) (Fig. 1e). In contrast, SARS2-N501Y_MA30_-infected mice failed to show a significant preference for the social stimulus at 20, 30-40, and 100 dpi (Fig. 1e). Social preference indices^47^ were calculated accordingly and found to be significantly decreased at 20 dpi and trend to significant reduction at 30-40 dpi compared with mock-infected controls (Fig. 1f), indicating persisting social interaction deficits. We also performed marble burying tests to evaluate changes in anxiety-like behaviours (Fig. 1g). Consistent with a previous study in hamsters^48^, infected mice showed significant reductions in burying activity at 30-40 dpi relative to controls (Fig. 1h), suggestive of an abnormal behavioural pattern^49, 50^. Collectively, these findings showed that peripheral sublethal infection with SARS2-N501Y_MA30_ induced various long-term behavioural changes in mice, in the apparent absence of virus neuroinvasion.

### Altered brain immune cell composition following peripheral SARS2-N501Y_MA30_ infection

While the mechanisms underlying neuroPASC are almost certainly multifactorial^22^, neuroinflammation is considered to be one of the major contributory factors in its development. Therefore, we next investigated the indirect effects of peripheral SARS2-N501Y_MA30_ infection of respiratory tract, sustentacular^43, 44^ or possibly other cells on immune cell responses in the brain. To this end, we isolated CD45^+^ immune cells from whole brains of SARS2-N501Y_MA30_-infected mice following perfusion at 6, 30, and 100 dpi, corresponding to acute, subacute, and long-term stages, respectively, and performed scRNA-seq (Fig. 2a). Mock-infected mice were processed in parallel as controls. This approach enabled us to track dynamic changes in brain immune cell composition and transcriptional profiles across disease stages. As a result, a total of 20,153 cells were recovered, revealing 11 immune cell types, including microglia, two discrete subsets of BAMs, monocytes, neutrophils, T and B lymphocytes, and minor populations of other immune cells (Fig. 2b, S1a-c and Table S1). The most prominent compositional shift was seen at 6 dpi, characterized by marked increases in monocyte and neutrophil frequencies compared with mock-infected controls (Fig. 2c).

**Figure 2.**
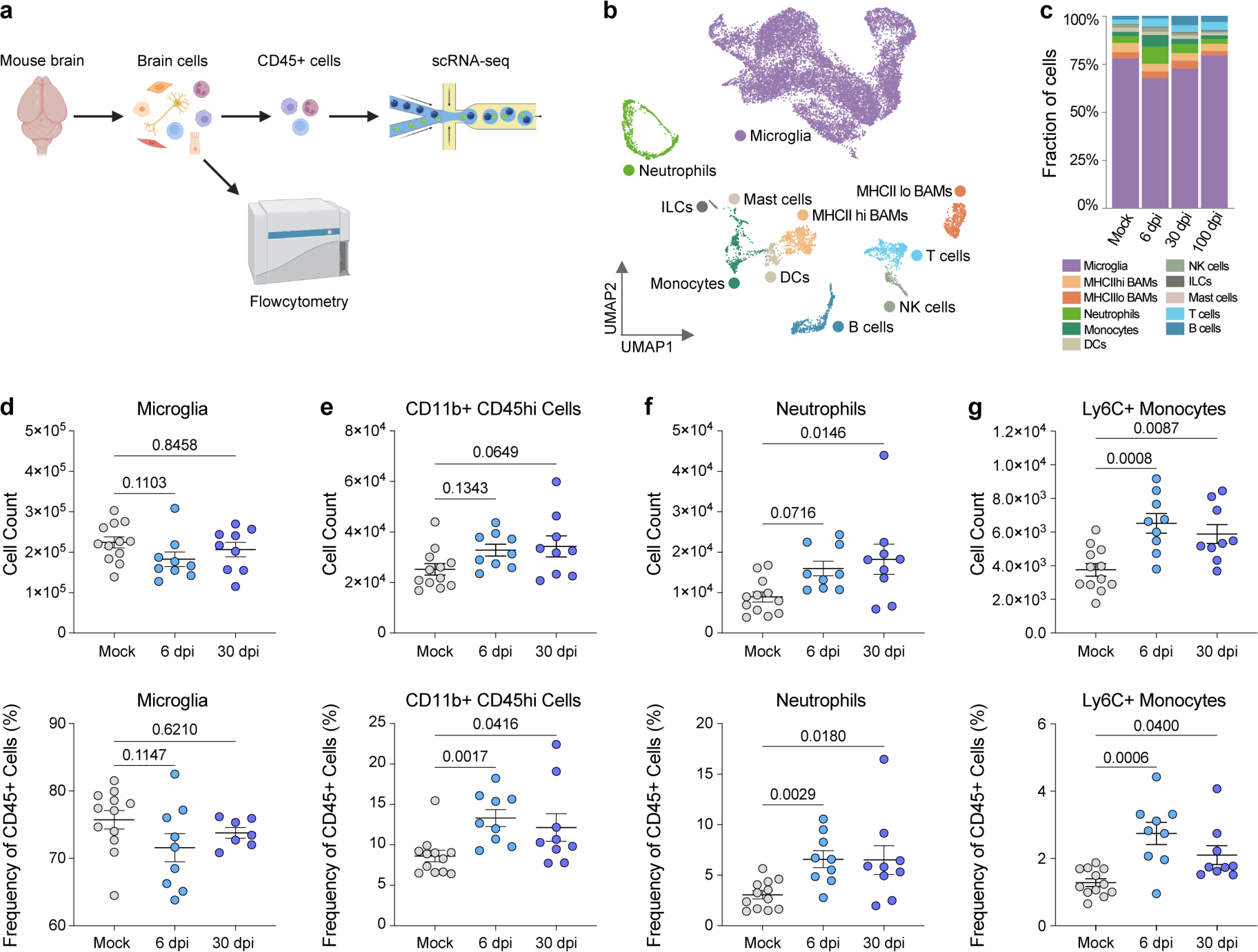
Peripheral myeloid cell infiltration into the brain following intranasal SARS2-N501Y_MA30_ infection. **a** | Schematic of the experimental workflow. Whole brains were dissociated, followed by either magnetic bead enrichment of CD45^+^ cells and single-cell RNA sequencing (scRNA-seq) or multiplex flow cytometry. **b** | Uniform Manifold Approximation and Projection (UMAP) visualization of brain immune cells identified by scRNA-seq, including samples from mock, 6, 30, and 100 days post-infection (dpi). BAMs, border-associated macrophages; DCs, dendritic cells; NK cells, natural killer cells; ILCs, innate lymphoid cells. **c** | Relative composition of brain immune cells across post-infection time points in the scRNA-seq dataset. **d–g** | Numbers and ratios of microglia, CD11b^+^CD45^hi^ cells, neutrophils, and Ly6C^+^ monocytes in brains at 6 and 30 dpi, measured by multiplex flow cytometry. Data in (**d–g**) are represented as mean with SEM and are pooled from two independent experiments per time point (mock: n = 12; 6 dpi: n = 9; 30 dpi: n = 9). Outliers were removed using the ROUT method (Q = 1%). Normality was assessed by the Shapiro–Wilk test; normally distributed data were compared by ordinary one-way ANOVA with Dunnett’s multiple comparisons test, while non-normally distributed data were analyzed using the Kruskal–Wallis test with Dunnett’s multiple comparisons test.

We subsequently validated these changes in cellular composition using multiplex flow cytometry. Myeloid cell populations were distinguished based on CD45 and CD11b expression (Fig. S2a). The number and frequency of microglia (CD45^mid^CD11b^+^ cells) did not significantly change between mock-infected and post-infection samples (Fig. 2d). In contrast, the frequency of CD45^hi^CD11b^+^ cells was elevated at both 6 and 30 dpi (Fig. 2e). Further analysis of CD45^hi^CD11b^+^ cells revealed increased infiltration of neutrophils (Ly6G^+^ cells) (Fig. 2f) and classical monocytes (Ly6C^+^Ly6G^-^ cells) (Fig. 2g). These results are generally consistent with the scRNA-seq results (Fig. 2c), although the magnitude of infiltration differed modestly when results from the two approaches were compared. On the other hand, Ly6C^-^Ly6G^-^ populations, corresponding to non-classical monocytes and BAMs, were stably populated across samples (Fig. S2b). Beyond myeloid cells, we also detected increased numbers and frequencies of B cells at 30 dpi and Ly6C^+^CD8^+^ T cells^51^ at 6 and 30 dpi but not total CD4 or CD8 T cells at either time point (Fig. S2c-f). Altogether, the presence of these infiltrating immune cells was consistent with a disturbance of brain immune cell homeostasis following pulmonary SARS2-N501Y_MA30_ infection, motivating subsequent transcriptional analyses of individual brain immune cell types.

### Microglial state heterogeneity and compositional shift after SARS2-N501Y_MA30_ infection

Microglia are considered to be key drivers of neuroinflammation. To delineate their contributions in SARS2-N501Y_MA30_-infected mice, we extracted all microglial cells (n = 15,086) from our scRNA-seq dataset and performed high-resolution subclustering (resolution = 2) to capture rare populations. Similar subclusters were merged based on their proximity in the UMAP space (Fig. S3a), shared marker gene expression, and inter-subcluster correlation (Fig. S3b), resulting in a final set of 9 subclusters (Fig. 3a, b and Table S2). A clear segregation between naïve and post-infection microglia was revealed by UMAP projection (Fig. 3c) and was accompanied by markedly different subcluster compositions (Fig. 3d), indicating broad microglial transcriptional alterations following SARS2-N501Y_MA30_ infection.

**Figure 3.**
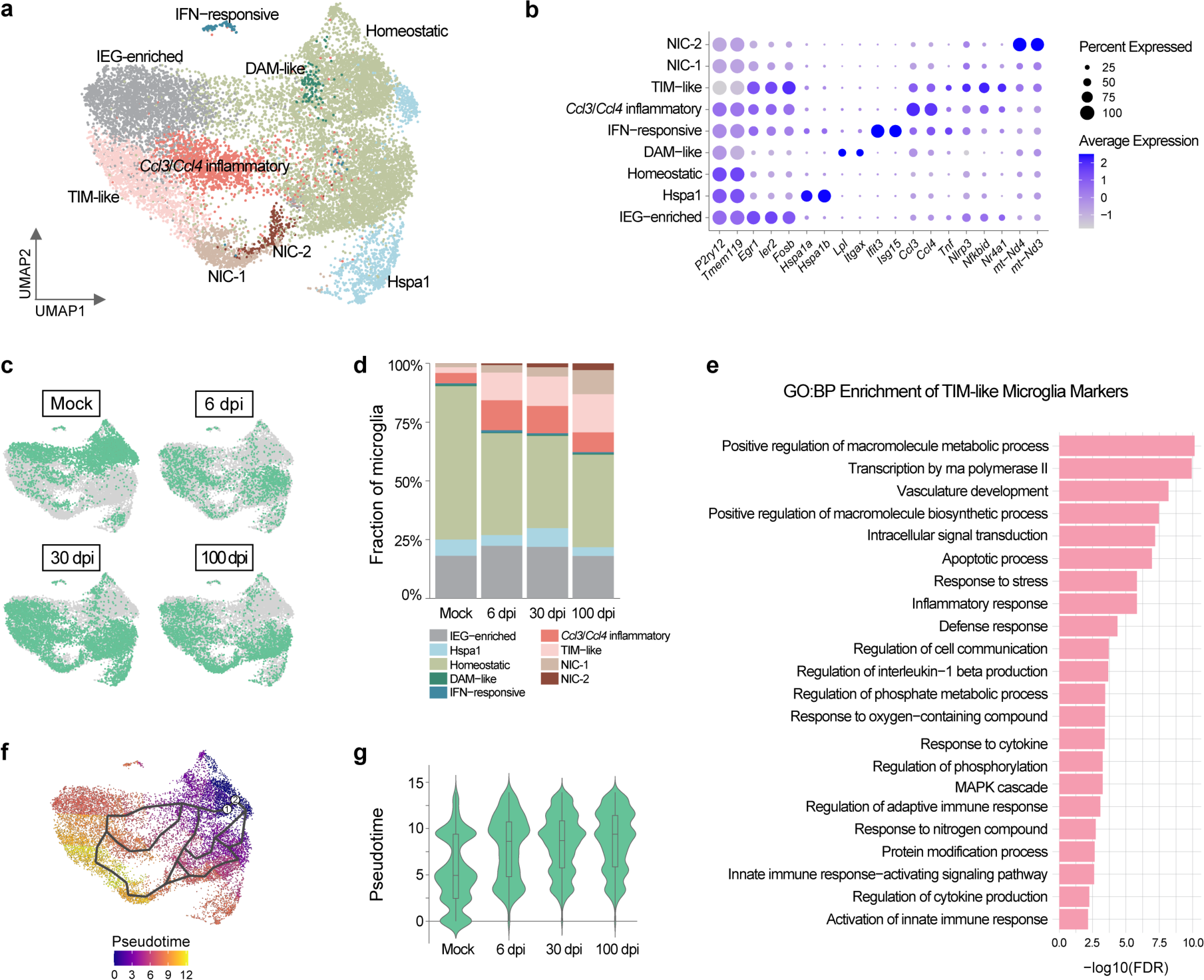
Microglia heterogeneity and long-term compositional shifts in response to intranasal SARS2-N501Y_MA30_ infection. **a** | UMAP plot showing subclustering of microglia from mock, 6, 30, and 100 dpi samples. IEG, immediate early gene; DAM, disease-associated microglia; TIM, terminally inflammatory microglia; NIC, non-inflammatory cluster. **b** | Dot plot showing representative marker genes for each microglial subcluster. **c** | UMAP plots illustrating sample-specific distributions of microglia. Cells belonging to the indicated sample are highlighted in green, whereas cells from all other samples are shown in grey. **d** | Relative proportions of microglial subclusters across time points. **e** | Gene ontology biological process (GO:BP) enrichment of TIM-like microglial signature genes. **f** | Pseudotime trajectory of microglial subclusters projected onto the UMAP. Homeostatic microglia from the mock-infected control were positioned at the root, with divergent branches capturing distinct activation pathways. **g** | Comparison of pseudotime distributions between samples.

Among identified clusters, one was present across all time points at comparable frequencies (Fig. 3d) and was primarily characterized by high expression of immediate-early genes (IEGs), including *Egr1* and *Ier2* (Fig. 3b). This pattern likely reflects dissociation-induced activation during tissue preparation^52, 53, 54^. We will resolve this in the future by adding the transcription inhibitor actinomycin D at the time of cell harvest to inhibit de novo RNA transcription^55^. Similarly, a smaller cluster was enriched for *Hspa1b* and *Hspa1a*, which have also been associated with dissociation-induced activation^52, 53, 54^. Beyond these two clusters, we identified a homeostatic subset abundantly present in all samples (Fig. 3a, d), defined by the high expression of canonical microglial homeostatic genes such as *P2ry12* and *Tmem119*, and *P2ry13*^56^ (Fig. 3b, S4a). However, homeostatic cells in mock-infected and post-infection samples showed minimal overlap in the UMAP space (Fig. 3c). Comparative analysis revealed upregulation of genes involved in the regulation of metabolic process and macromolecule modification in post-infection homeostatic microglia (Table S3). In addition, minor but disease-relevant populations were also detected. A small cluster (0.4-0.8% per sample) recapitulated disease-associated microglia (DAM)-like signatures, including upregulation of *Lpl* and *Itgax*^57^ (Fig. 3b). These cells are predicted to possess enhanced phagocytic activity compared to other microglia^58^. Unexpectedly, the frequency of DAM-like microglia was similar across samples. Additionally, interferon-responsive microglia were also detected at a low frequency, defined by elevated expression of interferon-stimulated genes such as *Ifit3* and *Isg15*^59^ (Fig. 3b). Notably, their frequency increased two-fold at 6 dpi before declining at 30 and 100 dpi, indicative of a transient antiviral response.

These clusters comprised over 90% of mock microglia. Several other clusters, on the other hand, were detected at low frequencies under mock-infected conditions but markedly expanded after infection. One such cluster was distinguished by high expression of inflammatory chemokines *Ccl3* and *Ccl4* (Fig. 3b), whose upregulation has been described in animal models of aging^60, 61^, brain injury^61^, and neurodegenerative diseases^60, 62^. These chemokines may function by regulating the trafficking and effector functions of peripheral immune cells^63^ or by directly influencing neuronal activity and survival^62^. This cluster also showed enrichment of *Ccrl2* and *Csf1* (Table S2), genes implicated in modulating microglial activation^64^ and in supporting microglial development and maintenance^65^, respectively. Another inflammatory cluster was annotated as terminally inflammatory microglia (TIM)-like, defined by concomitant expression of inflammatory genes (such as *Tnf* and *Nlrp3*) and stress markers (including *Egr1*, *Ier2*, and multiple AP-1 family transcription factors)^66^ (Fig. 3b). Gene Ontology (GO) enrichment analysis of TIM-like markers confirmed heightened activity of these cells in inflammatory and stress responses (Fig. 3e). Originally hypothesized to represent an exhausted-like state of inflammatory microglia in Alzheimer’s disease (AD)^66^, TIM-like cells in our dataset may represent a similar phenotype. We also identified two closely related non-inflammatory clusters (NICs) (Fig. 3b, S3c), both exhibiting high module scores for genes involved in RNA processing and protein turnover (Fig. S4c). One of these clusters lacked other clearly distinguishable features (Table S2, S3) and was designated NIC-1. The other cluster (NIC-2) showed elevated expression of mitochondrial DNA (mtDNA)-encoded transcripts (Fig. 3b, S4c) and was therefore termed high-mito microglia. Compared with homeostatic microglia, these four infection-enriched clusters showed reduced expression of homeostatic genes (Fig. S4a). Moreover, transcriptional profiles of TIM-like, NIC-1, and NIC-2 microglia correlated poorly with those of homeostatic microglia (Fig. S3c), highlighting their transcriptional divergence.

Pseudotime trajectory analysis using monocle3^67^ further reconstructed a continuum of microglial states. Homeostatic microglia from mock-infected mice were positioned at the root, with divergent branches capturing distinct activation pathways. TIM-like microglia occupied the most advanced pseudotime states (Fig. 3f), consistent with their designation as exhausted-like inflammatory cells. Overall, microglial pseudotime positions changed markedly over time compared with the mock-infected control (Fig. 3g), underscoring infection-driven transcriptional reprogramming.

### Persistent pro-inflammatory signatures and mitochondrial dysfunction in post-infection microglia

Microglial responses to SARS2-N501Y_MA30_ infection were further assessed by examining gene expression dynamics over time. Expression of homeostatic markers *P2ry12*, *Tmem11S*, and *P2ry13* was decreased in post-infection samples (Fig. S4b), indicating overall elevated microglial reactivity. Functional enrichment analysis of upregulated genes (avg_log2FC > 1, pct.1 > 0.2, and p_val_adj < 0.01) at 6 dpi revealed significant alterations in the regulation of immune responses (Fig. 4a). Notably, several immune-related changes persisted into the subacute and long-term phases. For example, *Tap1* and *H2-D1*, genes associated with MHC class I antigen presentation^68, 69^, along with additional genes in the KEGG (Kyoto Encyclopedia of Genes and Genomes) antigen processing and presentation pathway, remained consistently elevated at 6, 30, and 100 dpi relative to mock-infected samples (Fig. 4b). A group of inflammatory genes also exhibited sustained upregulation throughout these time points (Fig. 4c). In particular, although not uniformly produced in all microglia, elevated expression of proinflammatory cytokines – including *Tnf*, *Il1a*, *Il1b*, and *Il18*^70^ – was seen in all infected samples (Fig. 4c, d). In line with this, genes encoding key inflammasome components required for IL-1β and IL-18 maturation and release, including *Nlrp3*, *Nlrp1b*, *Casp1*, and *Gsdmd*^71, 72, 73^, were concordantly upregulated (Fig. 4c, d). Regulatory factors such as the nuclear receptor *Nr4a1* ^74^ and NF-κB inhibitors *Nfkbia* and *Nfkbid* ^75^ were also transcribed at higher levels in post-infection samples (Fig. 4c), suggesting a dynamic regulation of microglial inflammation.

**Figure 4.**
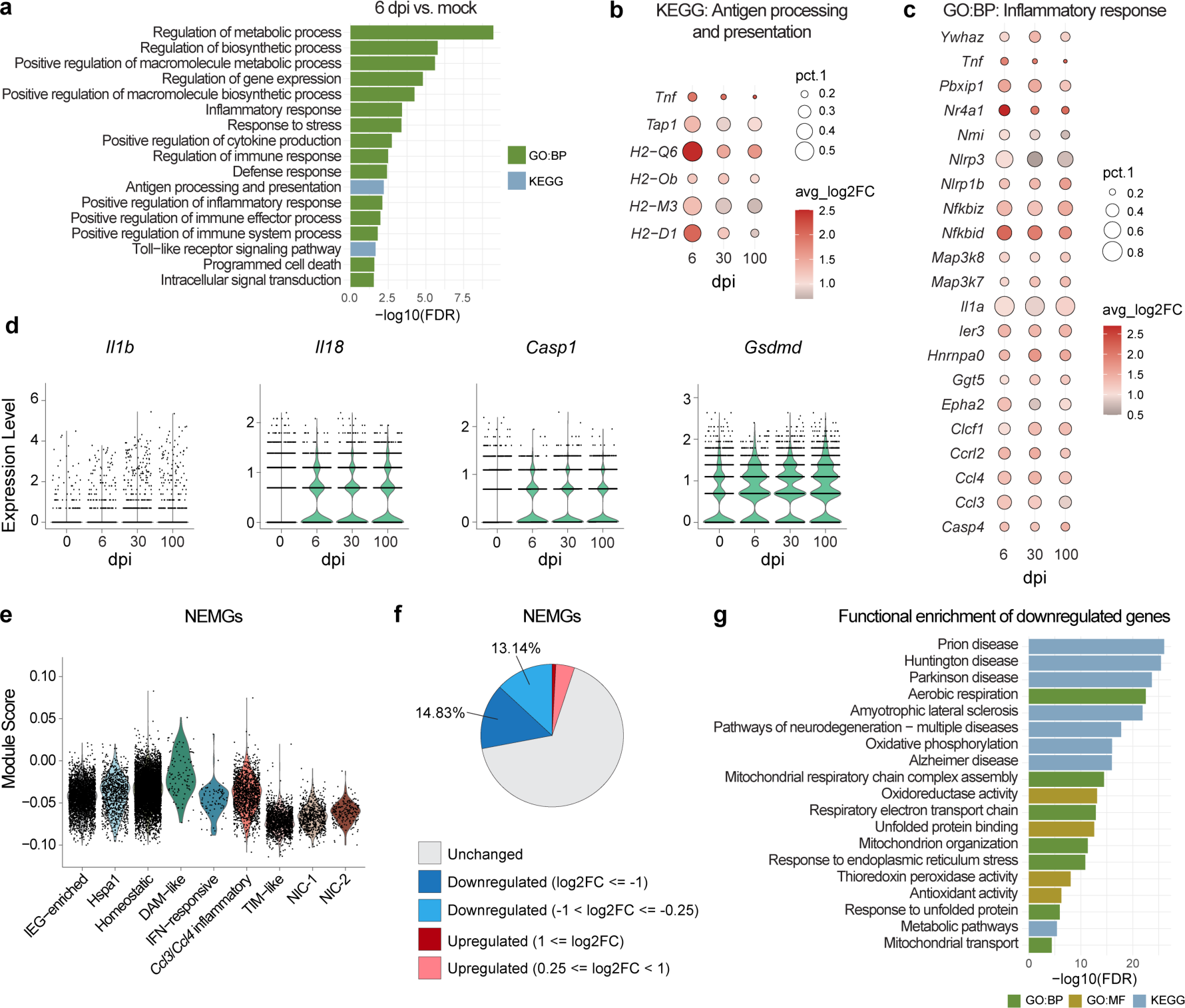
Persistent inflammatory response and mitochondrial dysfunction in post-infection microglia. **a** | Functional enrichment of genes upregulated at 6 dpi relative to mock. Selected terms are shown. KEGG, *Kyoto Encyclopedia of Genes and Genomes.* **b** | Regulation of genes across samples under the KEGG pathway *antigen processing and presentation*. *pct.1* indicates the proportion of microglia in each sample expressing indicated genes; *avg_log2FC* indicates the log2 fold change relative to mock. **c** | Regulation of genes across samples under the GO:BP term *inflammatory responses*. **d** | Expression levels of inflammasome-dependent cytokines and inflammasome components across samples. **e** | Module scores of nuclear-encoded mitochondrial genes (NEMGs) across microglial subclusters. **f** | Proportion of dysregulated NEMGs among total NEMGs in TIM-like, NIC-1, and NIC-2 subclusters compared with others. A cutoff of adjusted *p* < 0.01 was applied. **g** | Functional enrichment of downregulated genes in TIM-like, NIC-1, and NIC-2 subclusters compared with others. GO:MF, Gene Ontology Molecular Function.

Microglial metabolic failure and mitochondrial injury have been reported in post-mortem brain samples from COVID-19 patients ^12^. Consistent with this observation, our functional enrichment analysis of differentially expressed genes (DEGs) suggested broad metabolic changes in microglia across multiple post-infection time points (Table S3). Intriguingly, three infection-enriched subclusters (TIM-like, NIC-1, and NIC-2) displayed pronounced mitochondrial gene dysregulation, evidenced by markedly reduced module scores of nuclear-encoded mitochondrial genes (NEMGs)^76^ (Fig. 4e). Of 944 detectable NEMGs, 14.8% were significantly downregulated (log2FC ≤ –1) in these clusters (Fig. 4f), including 44 of 89 genes encoding oxidative phosphorylation (OXPHOS) subunits. This prominent downregulation of NEMGs was predicted to impair crucial cellular processes such as mitochondrial organization, OXPHOS, aerobic respiration, ER stress response, unfolded protein response, and antioxidation, and contribute to pathways implicated in neurodegenerative diseases, including prion disease, Huntington’s Disease, Parkinson’s Disease, and AD (Fig. 4g). Conversely, these clusters showed increased expression of mtDNA-encoded transcripts, particularly in NIC-2 microglia (Fig. S4c). Previous studies have demonstrated asynchronous kinetics between NEMGs and mtDNA-encoded genes^77^, and also shown that OXPHOS transcripts encoded by the two genomes increase in parallel but not concordantly^78^. However, the divergent expression we observed was unexpected and underscored mitochondrial dysfunction as a key feature of microglial responses to SARS2-N501Y_MA30_ infection.

### Post-infection monocyte recruitment and chemokine activity revealed by BAM transcriptomic changes

BAMs are another group of brain-resident myeloid cells located in the perivascular compartment, meninges, and CP. In our scRNA-seq dataset, we identified two clusters of cells expressing universal BAM signature genes, including *Mrc1*, *Pf4*, and *Ms4a7*^53, 79^ (Fig. 2b, S5a, and Table S1). They corresponded to two discrete populations: MHCII^lo^ and MHCII^hi^ BAMs (Fig. 2b). Unlike the dynamically shifting microglia composition described thus far (Fig. 3d), these BAM clusters showed stable relative abundance across different time points (Fig. 2c). Phenotypically, MHCII^lo^ BAMs exhibited high expression of M2 macrophage markers (*Mrc1* and *Cd163*)^80^ and multiple anti-inflammatory genes (*Gas6*, *Stab1*, and *Maf*), while MHCII^hi^ cells were enriched for antigen processing and presentation genes (*H2-Ab1*, *H2-Aa*, and *Cd74*) as well as proinflammatory genes (*Il1b*, *Icam1*, and *Fcgr4*) (Fig. 5a). These two populations also expressed distinct sets of chemokine and chemokine receptors (Fig. 5a), indicating complementary roles in chemotaxis. Based on reported tissue-specific signature genes^53, 81^, we expected that MHCII^lo^ BAMs originated primarily from the meningeal and perivascular niches, whereas the MHCII^hi^ population included meningeal, perivascular and CP BAMs (Fig. 5b).

**Figure 5.**
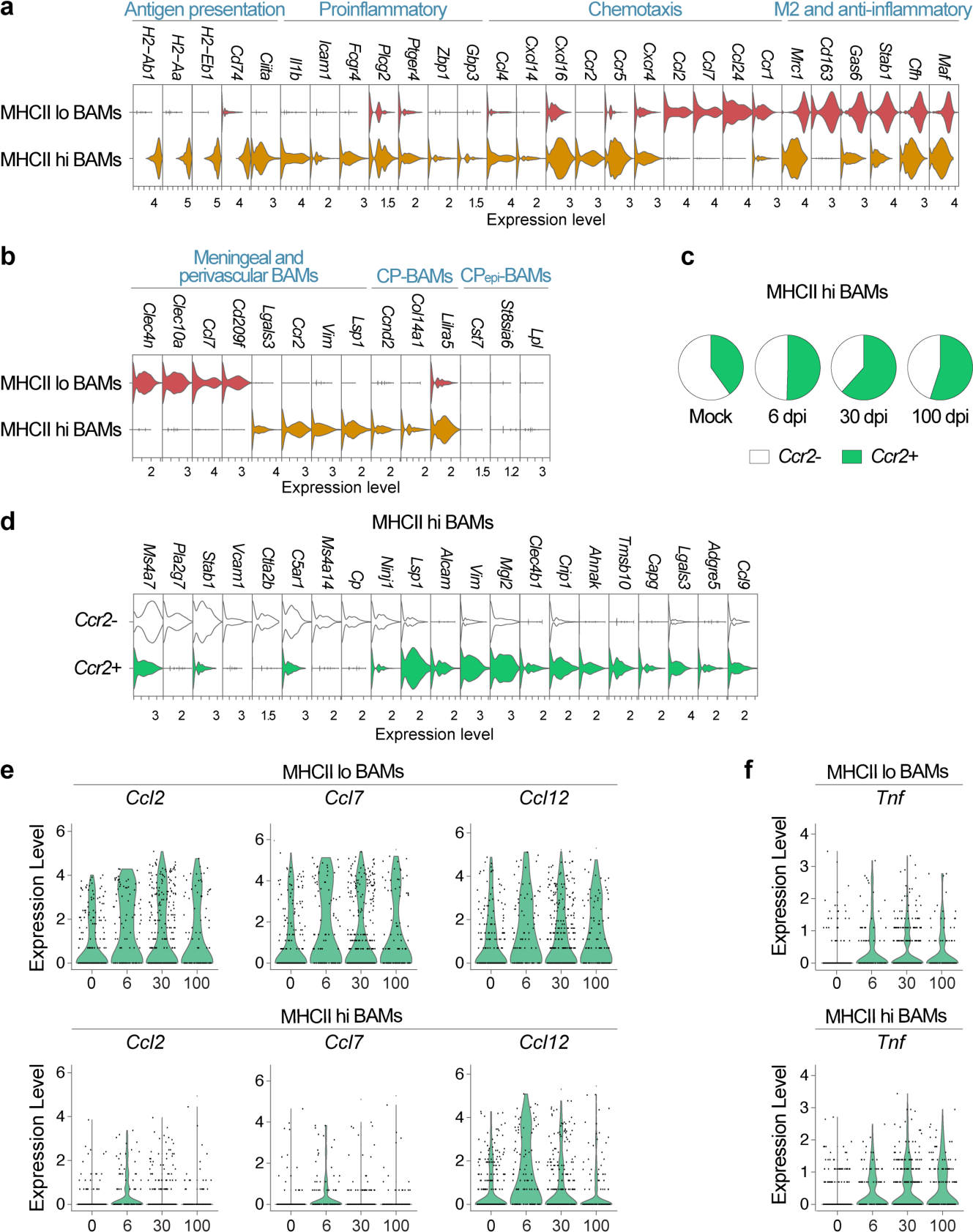
Monocyte recruitment and transient enhancement of chemokine activity in BAMs after SARS2-N501Y_MA30_ infection. **a** | Signature genes distinguishing MHCII^lo^ and MHCII^hi^ BAMs. **b** | Expression of tissue-specific signature genes in MHCII^lo^ and MHCII^hi^ BAMs. CP, choroid plexus. **c** | Proportion of *Ccr2*^+^ cells within MHCII^hi^ BAMs across samples. **d** | Signature genes distinguishing *Ccr2*^-^ and *Ccr2*^+^ MHCII^hi^ BAMs. **e** | Expression of the monocyte-attracting chemokines *Ccl2*, *Ccl7*, and *Ccl12* across samples. **f** | Expression of *Tnf* across samples.

Notably, *Ccr2*, a marker of infiltrating monocytes, was exclusively detected in MHCII^hi^ macrophages (Fig. 4a, b), consistent with a monocytic origin for a fraction of these cells^53^. This gradual renewal of specific BAM subsets by peripheral cells happens under homeostatic conditions^53^, but disruption of CNS homeostasis significantly accelerates this replenishment^82^. To investigate the impact of SARS2-N501Y_MA30_ infection, we compared the frequency of *Ccr2*^+^ macrophages at different post-infection time points. We found that *Ccr2*^+^ BAMs were most abundant at 30 dpi, increasing from 40.07% of total MHCII^hi^ BAMs in the mock control to 61.65% (Fig. 5c). This elevation suggested ongoing monocyte recruitment and transition to macrophages, reflecting peripheral infection– driven changes to brain homeostasis. By 100 dpi, the frequency of *Ccr2*^+^ macrophages declined, probably due to the gradual resolution of inflammation and phenotypic adaptation of infiltrating cells to the brain microenvironment niche. Indeed, the *Ccr2*^-^ population showed higher expression of certain BAM-associated genes, such as *Ms4a7*, *Stab1*, *Vcam1*, and *Cp*^53, 83^, while *Ccr2*^+^ cells maintained several monocyte-associated genes, including *Lsp1*, *Crip1*, *Tmsb10*, *Capg*, and *Adgre5*. Nevertheless, compared with *Ccr2*^+^ monocytes, *Ccr2*^+^ macrophages exhibited robust BAM signatures and high MHC II gene expression (Fig. S5b).

Consistent with the elevated frequency of *Ccr2*^+^ macrophages in post-infection samples, upregulation of monocyte-attracting chemokines, including *Ccl2*, *Ccl7*, and *Ccl12*^84^, was observed in both MHCII^lo^ and MHCII^hi^ BAMs at 6 dpi (Fig. 5e). Furthermore, the expression of *Tnf*, a prototypic proinflammatory cytokine, increased across all post-infection time points (Fig. 5f), indicating a persistently inflammatory milieu, potentially driven by infiltrating monocytes^85^. Supporting this, within the MHCII^hi^ BAM population, expression of *Il1b, Il18*, and their associated inflammasome components *Nlrp3*, *Casp1*, and *Gsdmd* was also upregulated (Fig. S5c).

### Inflammatory and antiviral programs in infiltrating monocytes and neutrophils

As shown by scRNA-seq and flow cytometry, monocyte and neutrophil frequencies in the brain markedly increased during acute infection (Fig. 2c, f, g). In mice, two major monocyte subsets have been described: classical Ly6C^+^ (CX3CR1^int^CCR2^+^CD62L^+^ CD43^low^Ly6C^hi^) and nonclassical Ly6C^-^ (CX3CR1^hi^CCR2^-^CD62L^-^CD43^hi^Ly6C^lo^)^86, 87^. Because *Ly6c2* transcripts were undetectable in our dataset for technical reasons^88^, we used *Ccr2* expression to distinguish these two populations. *Ccr2*^+^ cells constituted the majority of infiltrating monocytes, although *Ccr2*^-^ cells were also increased relative to the mock-infected control (Fig. 6a). Transcriptionally, *Ccr2*^+^ monocytes expressed higher levels of *Sell* (encoding CD62L) and *Lyz1*, consistent with a classical phenotype (Fig. 6b)^89^. Conversely, *Spn* (encoding CD43), *Itgax*, *Cd36*, *Pparg*, and *Fcgr4* were enriched in *Ccr2*^-^ cells, indicating a nonclassical signature^89, 90^. These findings align with the established role of classical monocytes as the primary subset recruited to inflamed tissues^86, 87^. DEG analysis between mock and infected samples underscored a strong antiviral response in monocytes at 6 dpi. Upregulated genes were functionally enriched in defense response to virus, cellular response to interferon-beta and type II interferon, activation of innate immune response, and negative regulation of viral process (Fig. 6c). Correspondingly, ISGs such as *Isg15*, *Oas2*, *Ifit3*, and *Irf7* were robustly induced at this time point (Fig. 6d). By 30 and 100 dpi, these genes and other antiviral signatures had returned to baseline levels (Fig. 6d). Moreover, apart from persistent *Il1b* upregulation (Fig. 6d), no sustained induction of inflammatory genes was detected (Fig. S6a).

**Figure 6.**
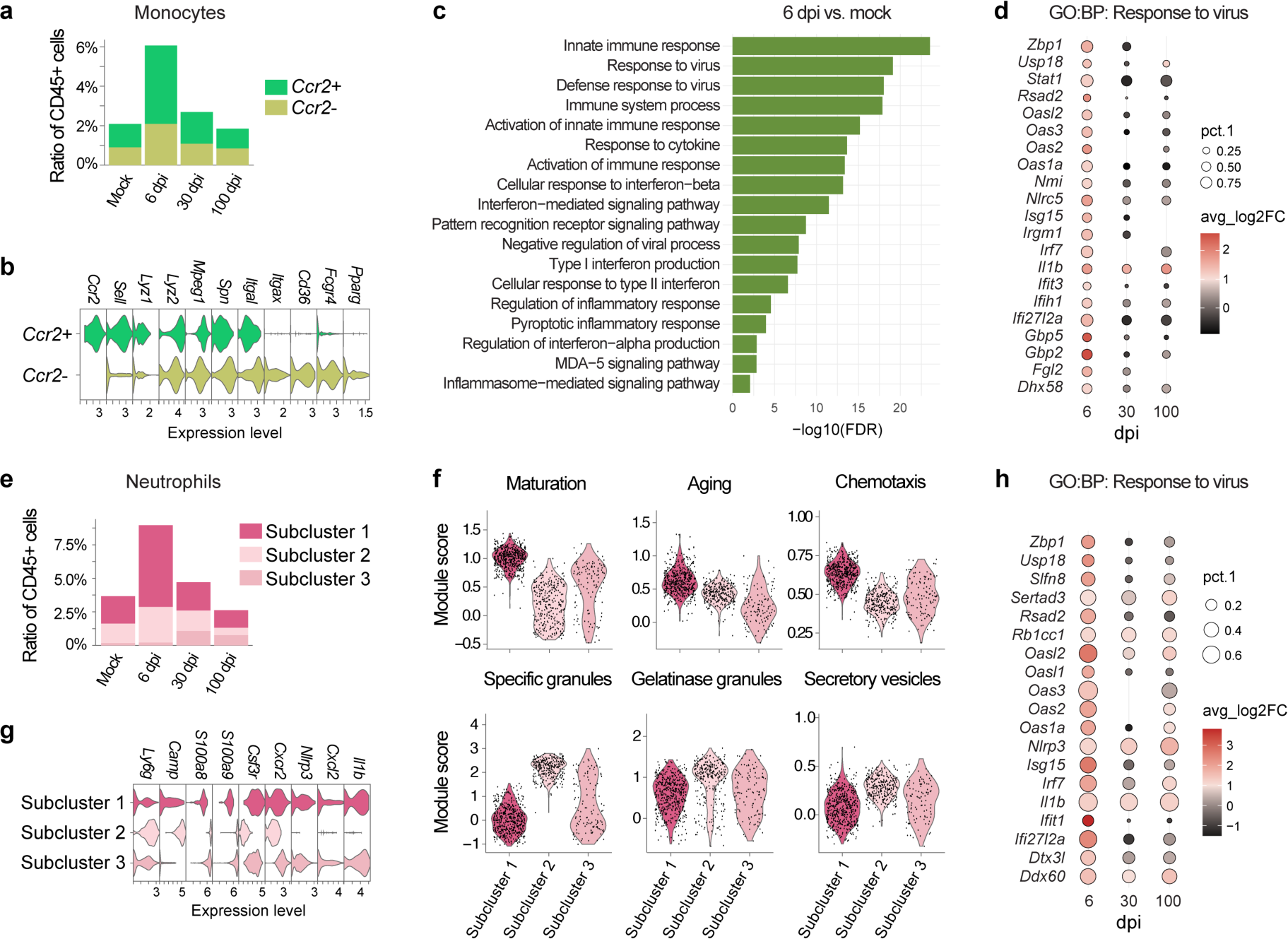
Acute infiltration of inflammatory monocytes and neutrophils into the brain following peripheral infection and their antiviral responses. **a** | Relative abundance of *Ccr2*^+^ and *Ccr2*^-^ monocytes across samples. **b** | Expression of classical and non-classical monocyte signature genes in *Ccr2*^+^ and *Ccr2*^-^ monocytes. **c** | GO:BP enrichment of genes upregulated in monocytes at 6 dpi relative to mock-infected samples. Selected terms are shown. **d** | Regulation of genes in monocytes across samples under the GO:BP term *response to virus. e |* Relative abundance of neutrophil subclusters across samples. **f** | Module scores of featured gene sets across neutrophil subclusters. **g** | Expression of neutrophil marker genes and typical inflammatory genes across neutrophil subclusters. **h** | Regulation of genes in neutrophils across samples under the GO:BP term *response to virus*.

Neutrophils were partitioned into three subpopulations via the initial unsupervised clustering (Fig. S1b). Their abundance varied across different time points, with subcluster 1 becoming prominently enriched at 6 dpi (Fig. 6e). This cluster displayed the highest maturation score (Fig. 6f), reflecting elevated expression of genes related to neutrophil development and terminal differentiation^91^. Neutrophils in this cluster appeared to be more “aged” than in other subclusters (Fig. 6f), a feature enabling these cells to more rapidly respond to inflammatory signals^92^. Moreover, subcluster 1 exhibited the highest chemotaxis score, but expressed various granule-associated genes at the lowest levels^91^ (Fig. 6f). In contrast, subcluster 2 showed a granule-enriched phenotype consistent with strong cytotoxic activity, whereas cluster 3 represented an intermediate state between subclusters 1 and 2 (Fig. 6f). Notably, subclusters 1 and 3 uniquely expressed inflammatory mediators such as *Il1b*, *Cxcl2*, and *Nlrp3* (Fig. 6g). Collectively, these findings indicated that brain-infiltrating neutrophils following SARS2-N501Y_MA30_ infection predominantly skewed toward a mature, aged, and inflammatory state (subcluster 1). Similar to monocytes, neutrophils also mounted a transient antiviral response at 6 dpi (Fig. 6h, S6b), highlighting their involvement in the early response to peripheral viral infection.

## DISCUSSION

Neuroinflammation is a prominent feature observed in COVID-19 patients with acute and persistent neurological abnormalities^10, 12, 13, 14, 15^. Here, using an established mouse model of neuroPASC, we described a comprehensive analysis of the longitudinal contributions of different brain myeloid cells to this process. We showed that microglia underwent profound and lasting transcriptional shifts after peripheral SARS2-N501Y_MA30_ infection (Fig. 3c, d, g). Most notably, expression of multiple inflammatory mediators was consistently upregulated across acute, subacute, and long-term time points (Fig. 4c, d), indicative of their major role in chronic neuroinflammation. Persistent elevation of pro-inflammatory cytokines was also observed in another brain-resident macrophage population, BAMs (Fig. 5f, S5b). Meanwhile, these cells transiently enhanced the production of monocyte attractants during acute infection (Fig. 5e), concordant with increased *Ccr2*^+^ cell frequency at later time points (Fig. 5c), suggesting disturbed brain homeostasis. In contrast to sustained alterations in resident myeloid cells, inflammatory monocytes and neutrophils were transiently recruited into the brain and exhibited strong antiviral signatures (Fig. 2, 6).

Our findings recapitulate several hallmarks of microglial reactivity previously reported in COVID-19 post-mortem brain samples. Similar to those studies, we observed the emergence of COVID-19-associated microglial clusters^14^ (Fig. 2c, d), downregulation of homeostatic markers such as *P2YR12*^12, 14, 15^ (*P2ry12* in mice) (Fig. S4b), and mitochondrial injury reflected by metabolic pathway dysregulation and significant suppression of mitochondrial gene expression^12^ (Fig. 4e-g). Importantly, our data revealed that these alterations were initiated during acute infection and persisted for at least 100 dpi, whereas existing human datasets largely represent acute or subacute disease stages. Moreover, in line with their established role in neuroinflammation, post-infection microglia exhibited sustained upregulation of neurotoxic factors such as *Tnf* and *Il1b* (Fig. 4c, d).

Meanwhile, some microglial features we identified are like those reported in neurodegenerative conditions. The TIM-like subcluster, which was almost exclusively enriched in post-infection samples (Fig. 3d), resembles an exhausted-like microglial population in brains of aged AD mice bearing the human *APOE4* allele, the single greatest monoallelic risk factor for late-onset AD^66^. In experimental autoimmune encephalomyelitis, a model of human MS, a subpopulation of DAMs in the spinal cord upregulates both nuclear and mtDNA-encoded genes for mitochondrial complex I (CI) subunits, sustaining microglial activation and neuroinflammation through CI-driven reverse electron transport and consequent reactive oxygen species production^93^. In parallel, the NIC-2 subcluster in our post-infection samples showed enhanced expression of all mtDNA-encoded CI subunit genes. Although augmented expression of nuclear-encoded CI components (e.g., *Ndufs4*) and oxidative stress markers (e.g., GP91-PHOX, *Sod1*) was not evident, the transcriptional profile of NIC-2 microglia may reflect mitochondrial dysfunction coupled to neuroinflammation. However, the core DAM signature, originally described in AD^57^ and later generalized to aging and diverse CNS diseases^58^, was only modestly enriched in a small fraction of microglia in our study (Fig. 3a-c, S4c). Since DAMs feature increased expression of genes involved in the processing and clearance of phagocytosis substrates^58^, peripheral SARS2-N501Y_MA30_ infection likely does not substantially enhance microglial phagocytic activity. Instead, the downregulation of homeostatic genes together with persistent inflammatory activation in post-infection microglia more closely resembles transcriptional changes induced by systemic LPS challenge^94^.

In addition to parenchymal microglia, our scRNA-seq data identified MHCII^hi^ and MHCII^lo^ BAMs and *Ccr2*^+^ and *Ccr2*^-^ monocytes in mouse brains. Their compositional and transcriptional shifts collectively underscored the recruitment of *Ccr2*^+^ monocytes and transition into MHCII^hi^ macrophages in response to SARS2-N501Y_MA30_ infection (Fig. 5c, 6a). While circulating monocytes minimally enter the healthy CNS, they infiltrate the brain in different CNS disease contexts and contribute to an inflammatory immune response^95, 96^, similar to our observations here. Notably, although inflammatory response genes were enriched in monocytes only at 6 dpi (Fig. S6a), persistent inflammation markers were found in BAMs (Fig. 5f), especially the MHCII^hi^ subpopulation (Fig. S3b). Moreover, the upregulation of monocyte attractants in BAMs (*Ccl2*, *Ccl7*, *Ccl12*) and microglia (*Ccl3* and *Ccl4*) at 6 dpi suggests that they may cooperatively promote monocyte recruitment into the CNS.

Although less well studied, a growing body of evidence has shown the contribution of neutrophils to CNS disease pathogenesis. Their extravasation into the brain and subsequent release of neutrophil extracellular traps (NETs) and IL-17 are reported in AD neuropathology^97^, where IL-17 interacts with microglia to suppress their response to neurodegeneration^98^. Other neurodegenerative diseases can also be exacerbated by neutrophils, producing an inflammatory state and neuronal death^99^. In SARS2-N501Y_MA30_ infected mice, inflammatory neutrophils, instead of those enriched in cytotoxic responses, migrated into the brain during acute infection (Fig. 2c, f, 6e-g), likely functioning through cell-cell communication with other immune cells.

Indeed, SARS2-N501Y_MA30_ infection appears to reshape the brain immune cell communication landscape, as inferred by CellChat, a computational tool analysing cell-cell communication networks from single-cell transcriptomic data^100^ (Fig. S7). Interactions between monocytes and microglia were shown to be enhanced at 6 dpi, followed by a decline at 30 dpi and back to mock-infected levels at 100 dpi. Neutrophil-microglia interactions were also stronger at 6 and 30 dpi. Notably, consistent with previous studies^10, 12^, CellChat analyses indicated elevated interaction strength between T cells and microglia. We also identified significant enrichment of genes related to T cell activation and differentiation across post-infection time points (Table S4). Meanwhile, enhanced B cell-microglia crosstalk occurred at 30 dpi, suggestive of a potentially delayed contribution of B cells to neuroPASC.

A limitation of the present study is that it primarily relies on scRNA-seq analysis. Despite its high resolution and breadth, there remains a significant gap in understanding how these transcriptional shifts relate to functional alterations^58^. For instance, while we identified marked downregulation of NEMGs in TIM-like, NIC-1, and NIC-2 microglia, the functional consequences of these metabolic disturbances remain to be further defined^101^. In addition, the Fixed RNA Profiling workflow used here restricts detection to a predefined probe set targeting the mouse transcriptome. As a result, key genes such as the classical monocyte marker gene *Ly6c2*, microglia marker *Cx3cr1*, and cytokines *Cxcl8* and *Il10* were undetectable or captured at extremely low levels, potentially obscuring certain immune traits relevant to neuroPASC pathogenesis.

Altogether, our study delineates the dynamic immune landscape underlying neuroPASC development. Monocytes and neutrophils exhibited strong antiviral signatures during acute infection, likely reflecting direct responses to peripheral viral challenge. Their subsequent infiltration into the CNS may initiate the persistent inflammation observed in microglia and BAMs, thereby contributing to a variety of neuroPASC symptoms. While we focused on only immune components here, the presence of marked leukocyte infiltration also supports the hypothesis that BBB disruption and endothelial inflammation following SARS2-N501Y_MA30_ infection contribute to the inflammatory milieu. Further work will be required to investigate the specific downstream impact of microglia inflammation on neuronal integrity. Potential consequences include accelerated accumulation of senescent cells leading to premature brain aging and neurodegeneration^102^, loss of hippocampal neurogenesis associated with cognitive deficits^8, 29^, and neurotransmitter dysregulation leading to corresponding neurological symptoms^15, 45^. Together, these findings provide a framework for understanding the immunopathological basis of neuroPASC and highlight potential avenues for therapeutic interventions such as limiting peripheral myeloid cell infiltration during acute infection and mitigating chronic microglia reactivity.

## MATERIALS AND METHODS

### Mice and viruses

4-6-month-old C57BL/6 mice (Charles River Laboratories) were used in all experiments. Mice were housed in the Animal Care Unit at the University of Iowa under standard conditions of dark/light cycle, ambient temperature and humidity. For each experiment, mouse ages were carefully matched, and animals were randomly assigned to experimental groups, with group sizes sufficient to achieve statistical significance.

SARS2-N501Y_MA30_ was propagated as previously described^43^. All virus stocks were sequenced following propagation and confirmed to match the input strain.

### Mouse infection

Mice were anaesthetized with a ketamine–xylazine cocktail (87.5 mg/kg ketamine and 12.5 mg/kg xylazine) and infected intranasally with 1000 PFU of virus in a total volume of 50 μL DMEM. Animal weight and health were monitored daily. All experiments involving SARS2-N501Y_MA30_ were conducted in either biosafety level 3 (BSL-3) or BSL-2 facilities at the University of Iowa. All animal studies were approved by the University of Iowa Animal Care and Use Committee and complied with the Guide for the Care and Use of Laboratory Animals.

### RNA isolation and RT-qPCR

Total RNA was extracted from tissues using TRIzol (Invitrogen) according to the manufacturer’s protocol. For first-strand cDNA synthesis, 3 μg of total RNA was used as template with M-MLV reverse transcriptase (Invitrogen). The resulting cDNA was subjected to amplification of selected genes by real-time quantitative PCR using Power SYBR Green PCR Master Mix (Applied Biosystems). Average values from technical duplicates were used to calculate relative transcript abundance, normalized to *Hprt1*, and presented as 2^−ΔCT^. The following primers were used: SARS-CoV-2 genome: F: 5′-GACCCCAAAATCAGCGAAAT-3′; R: 5′-TCTGGTTACTGCCAGTTGAATCTG-3′. *Hprt1*: F: 5′-GCGTCGTGATTAGCGATGATG-3′; R: 5′-CTCGAGCAAGTCTTTCAGTCC-3′.

### Mouse behavioral testing

For the sociality preference test, a custom-built three-chamber plastic apparatus (60 cm × 40 cm × 20 cm) with open doors connecting each chamber was used. Prior to testing, two clean empty cups were placed in side chambers, and the test mouse was allowed to habituate to the apparatus and empty cups for 5 min. Subsequently, an age-, sex-, and strain-matched unfamiliar mouse was placed under one cup to serve as the social stimulus, while a toy was placed under the other cup to serve as the non-social stimulus. The test mouse was then placed in the centre chamber and allowed to explore freely for 5 min. Time spent interacting with the social stimulus (T_S_) and the non-social stimulus (T_NS_) was recorded. Social preference indexes (I_SP_) were calculated as: I_SP_ = (T_S_ − T_NS_)/(T_S_ + T_NS_).

For the marble burying test, each test mouse was placed in a corner of a cage containing clean bedding with 20 equally spaced glass marbles arranged on the surface. Mice were allowed to explore freely for 15 min, after which numbers of buried and unburied marbles were recorded. A marble was considered buried if ≥60% of its surface was visually judged to be covered by bedding.

### Brain cell preparation and antibodies for flow cytometric analysis

Animals were deeply anaesthetized with ketamine–xylazine and perfused transcardially with 10 mL PBS. Intravenous labeling was not performed due to technical difficulties in the BSL3 lab. In preliminary experiments, we found that the vast majority of brain-associated immune cells were in the brain and not in the vasculature. Brains were dissected and digested in HBSS buffer containing 2% fetal calf serum, 25 mM HEPES, 1 mg mL^−1^ collagenase D (Roche) and 0.1 mg ml^−1^ DNase I (Roche) at 37 °C for 30 min using a gentleMACS dissociator (Miltenyi). Single-cell suspensions were prepared by passing through a 70-μm cell strainer, followed by debris removal with the Debris Removal Solution (Miltenyi). Red blood cells were lysed using the 10X Red Blood Cell Lysis Solution (Miltenyi). Remaining cells were enumerated using a Countess 3 cell counter (Invitrogen). Dead cells were then labeled with the Fixable Blue Dead Cell Stain Kit (Invitrogen). Cells were washed, pelleted, and blocked with 1 μg anti-CD16/anti-CD32 (2.4G2) at 4 °C for 5 min, followed by surface staining with the following antibodies at 4 °C for 30 min: BV450 anti-mouse CD45 (clone 30-F11, BioLegend, catalogue number 103135); eFluor450 anti-mouse CD11b (clone M1/70, eBioscience, catalogue number 48-0112-82); FITC anti-mouse CD11c (clone N418, eBioscience, catalogue number 11-0114-82); PE/Cy7 anti-mouse Ly6G (clone RB6-8C5, abcam, catalogue number ab25514); PerCP-Cy5.5 anti-mouse Ly6C (clone HK1.4, eBioscience, catalogue number 45-5932-82); BV605 anti-mouse I-A/I-E (clone M5/114.15.2, BioLegend, catalogue number 107639); BV421 anti-mouse CD3 (clone 17A2, BioLegend, catalogue number 100227); BUV563 anti-mouse CD4 (clone GK1.5, BD Biosciences, catalogue number 612923); APC anti-mouse CD8a (clone 53-6.7, eBioscience, catalogue number 17-0081-82); BV785 anti-mouse CD19 (clone 6D5, BioLegend, catalogue number 115543); APC/Cy7 anti-mouse P2RY12 (clone S16007D, BioLegend, catalogue number 848023). Antibodies were diluted according to manufacturers’ instructions. After staining, cells were washed and fixed with Cytofix (BD Biosciences). Flow cytometry data were acquired on a Cytek Aurora spectral flow cytometer (Cytek Biosciences) using SpectroFlo software and analyzed using FlowJo software (FlowJo). The gating strategy is shown in Figure S2a.

### scRNA-seq

Animals were deeply anesthetized with ketamine–xylazine and perfused transcardially with 10 mL PBS. Brains were collected, and brain single-cell suspensions were prepared using the Adult Mouse Brain Dissociation Kit (Miltenyi) according to the manufacturer’s instructions. CD45^+^ cells were then enriched by staining with CD45 magnetic microbeads and applying them to an LS column (Miltenyi) following the manufacturer’s protocol. Isolated cells were pelleted and fixed overnight in 10x Genomics Fixation Buffer (10x Genomics). Samples from 4–5 mice per experimental time point were pooled before submission to the University of Illinois Urbana-Champaign Genomics Core for processing with the 10x Genomics Chromium Fixed RNA Profiling (Flex) Gene Expression assay (10x Genomics). All libraries were prepared and sequenced together on the Illumina NovaSeq X Plus System, generating 28 × 90 nt reads. Notably, probes targeting the SARS-CoV-2 ORF1ab and nucleocapsid (N) genes were custom-designed and spiked into the Chromium Mouse Transcriptome Probe Set (v1.0.1), following the manufacturer’s instructions. Probe sequences are as follows:

ORF1ab: TACTAGTGCCTGTGCCGCACGGTGTAAGACGGGCTGCACTTACACCGCAA; ATAGATTACCAGAAGCAGCGTGCATAGCAGGGTCAGCAGCATACACAAGT; GAGCAAGAACAAGTGAGGCCATAATTCTAAGCATGTTAGGCATGGCTCTA.

N: CCATTCTGGTTACTGCCAGTTGAATCTGAGGGTCCACCAAACGTAATGCG; AGTTGTAGCACGATTGCAGCATTGTTAGCAGGATTGCGGGTGCCAATGTG; TGGAACGCCTTGTCCTCGAGGGAATTTAAGGTCTTCCTTGCCATGTTGAG.

### scRNA-seq data processing and analysis

The quality of raw sequencing data was assessed using FastQC (v0.12.1), and summary reports were compiled with MultiQC (v1.12). Gene-count matrix was obtained by aligning reads to a customized reference genome consisting of the GRCm38 (mm10) assembly along with the SARS-CoV-2 sequence (EPI_ISL_1666328) using Cell Ranger (v7.2.0). Subsequent analysis was conducted in R (v4.5.0) using the Seurat package (v5.3.0). Cell-level quality control was applied using the following thresholds: 400 < number of genes < 8,000; 800 < number of counts < 35,000; proportion of reads with mitochondrial origin < 5%^93^; log₁₀(number of genes) / log₁₀(number of counts) > 0.80. To focus on neuroimmune populations, non-immune cells were excluded based on the absence of *Ptprc* or *Itgam* expression using the subset function in Seurat. Genes detected in fewer than ten cells were also excluded from downstream analysis. The filtered data were normalized and scaled using SCTransform, with regression of cell cycle scores and mitochondrial gene content. Principal component analysis was conducted using default parameters. The first 40 principal components were used to compute a shared nearest neighbor graph and perform unsupervised clustering at a resolution of 0.6. To reduce ambient RNA artifacts, contamination in individual cells was estimated using DecontX (v1.6.0)^103^, with Seurat-derived clustering labels provided as priors. Clusters with high estimated contamination – confirmed by the expression of marker genes for endothelial cells, astrocytes, or neurons – were removed from downstream analysis. The DecontX-corrected count matrix was then reprocessed with SCTransform, followed by clustering at a resolution of 0.6. Clusters were manually annotated based on top upregulated genes identified by Seurat’s FindMarkers function, combined with reference to previously reported marker genes. Microglia were extracted from total cells using the subset function and further subclustered following SCTransform normalization and scaling.

Pseudotime trajectory analysis in microglia was performed with monocle3 (v1.4.26)^67^, with Seurat-derived UMAP embeddings and subcluster assignments used as inputs for trajectory graph learning and pseudotime measurement. Correlation matrices of microglial subclusters were generated from log-normalized gene expression values, and Pearson correlation coefficients were calculated. Module scores were calculated using the AddModuleScore function in Seurat, and genes included in each module are listed in Table S5. DEG analyses between conditions were conducted with the FindMarkers function in Seurat. Genes meeting the following thresholds were used for functional enrichment in gProfiler^104^: avg_log2FC >= 1, p_val_adj <= 0.01, pct.1 > 0.2 or avg_log2FC <= -1, p_val_adj <= 0.01, pct.2 > 0.2. Cell-cell communication analysis was conducted using the CellChat (v2.2.0) package^100^.

### Statistical analysis

Statistical analyses were performed using GraphPad Prism (v10.3.1) for vRNA RT–qPCR, behavioural test, and multiplex flow cytometry data. Two-way ANOVA with Sidak’s multiple comparisons test or one-way ANOVA with Dunnett’s multiple comparisons test was applied as appropriate.

### Data availability

Complete scRNA-seq data were deposited in the National Center for Biotechnology Information BioProject database under accession number XXXX (number pending).

