## supplemental_figures for "Longitudinal analysis reveals myeloid cell contributions to neuroPASC pathogenesis"

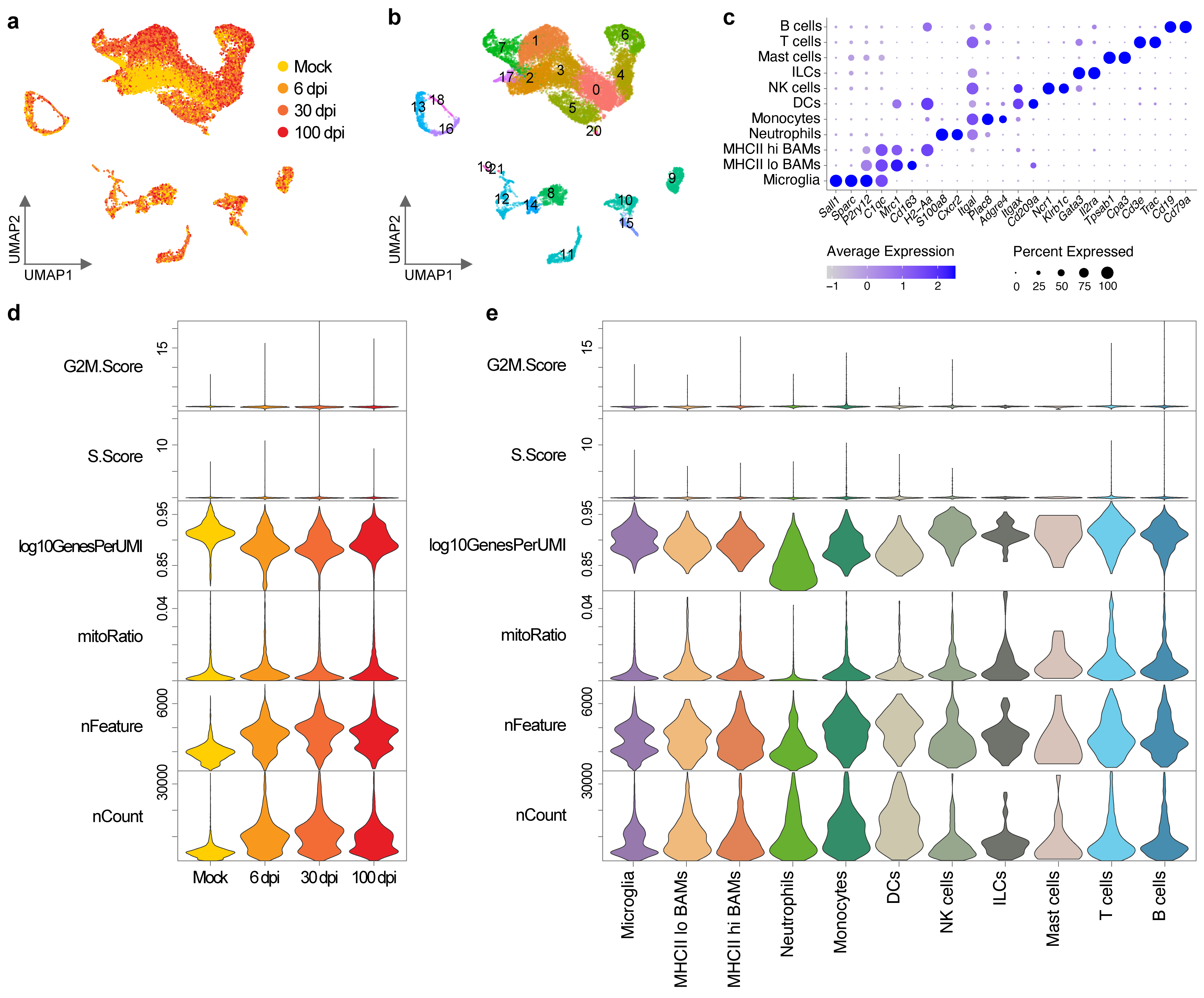

**Figure S1. Clustering and quality control (QC) of single-cell RNA sequencing (scRNA-seq) data.** **a** | Uniform Manifold Approximation and Projection (UMAP) plot showing the distribution of cells from mock-infected, 6, 30, and 100 days post-infection (dpi) samples. **b** | UMAP plot illustrating the initial clustering result prior to cell type annotation. **c** | Dot plot showing representative marker genes for each identified cell type. **d** | Violin plots showing QC metrics across samples. *G2M.Score*: module score for genes associated with G2/M cell cycle phases; *S.Score*: module score for genes associated with the S phase; *log10GenesPerUMI*: log₁₀(number of genes per cell / number of UMIs per cell), reflecting transcriptome complexity; *mitoRatio*: fraction of reads mapped to mitochondrial genes; *nFeature*: number of detected genes; *nCount*: total UMI counts per cell. **e** | Violin plots showing QC metrics across different cell types.

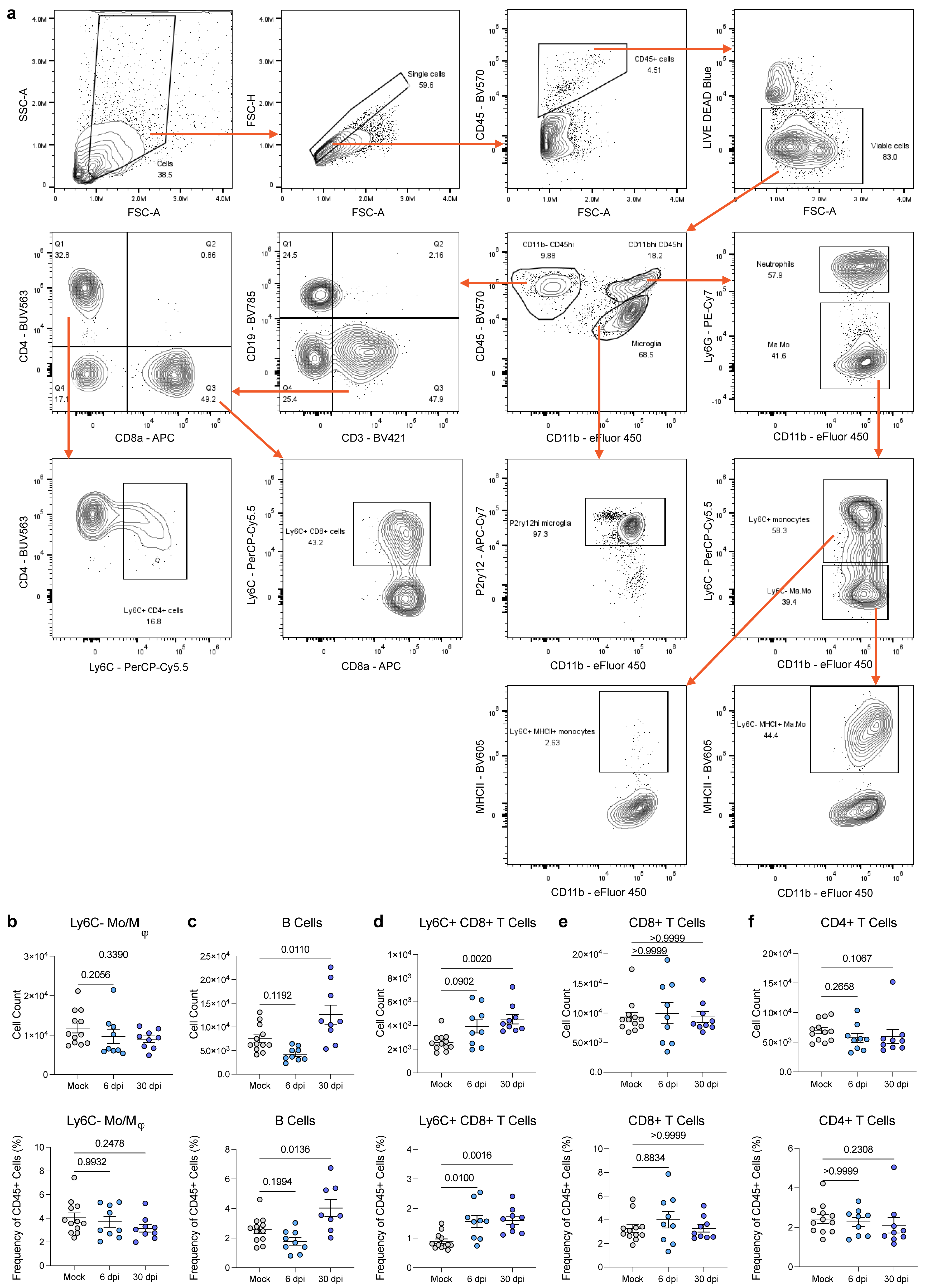

**Figure S2. Flow cytometric validation of brain immune cell composition changes following intranasal SARS2-N501Y_MA30_ infection.** **a** | Gating strategy of multiplex flow cytometry. **b–f** | Numbers and frequencies of Ly6C^-^ monocytes and macrophages, B cells, Ly6C^+^CD8^+^ T cells, CD8^+^ T cells, and CD4^+^ T cells in brains at 6 and 30 dpi. Data are represented as mean ± standard error of the mean (SEM) and are pooled from two independent experiments per time point (mock: n = 12; 6 dpi: n = 9; 30 dpi: n = 9). Outliers were removed using the ROUT method (Q = 1%). Normality was assessed by the Shapiro–Wilk test; normally distributed data were compared by ordinary one-way ANOVA with Dunnett’s multiple comparisons test, while non-normally distributed data were analyzed using the Kruskal–Wallis test with Dunnett’s multiple comparisons test.

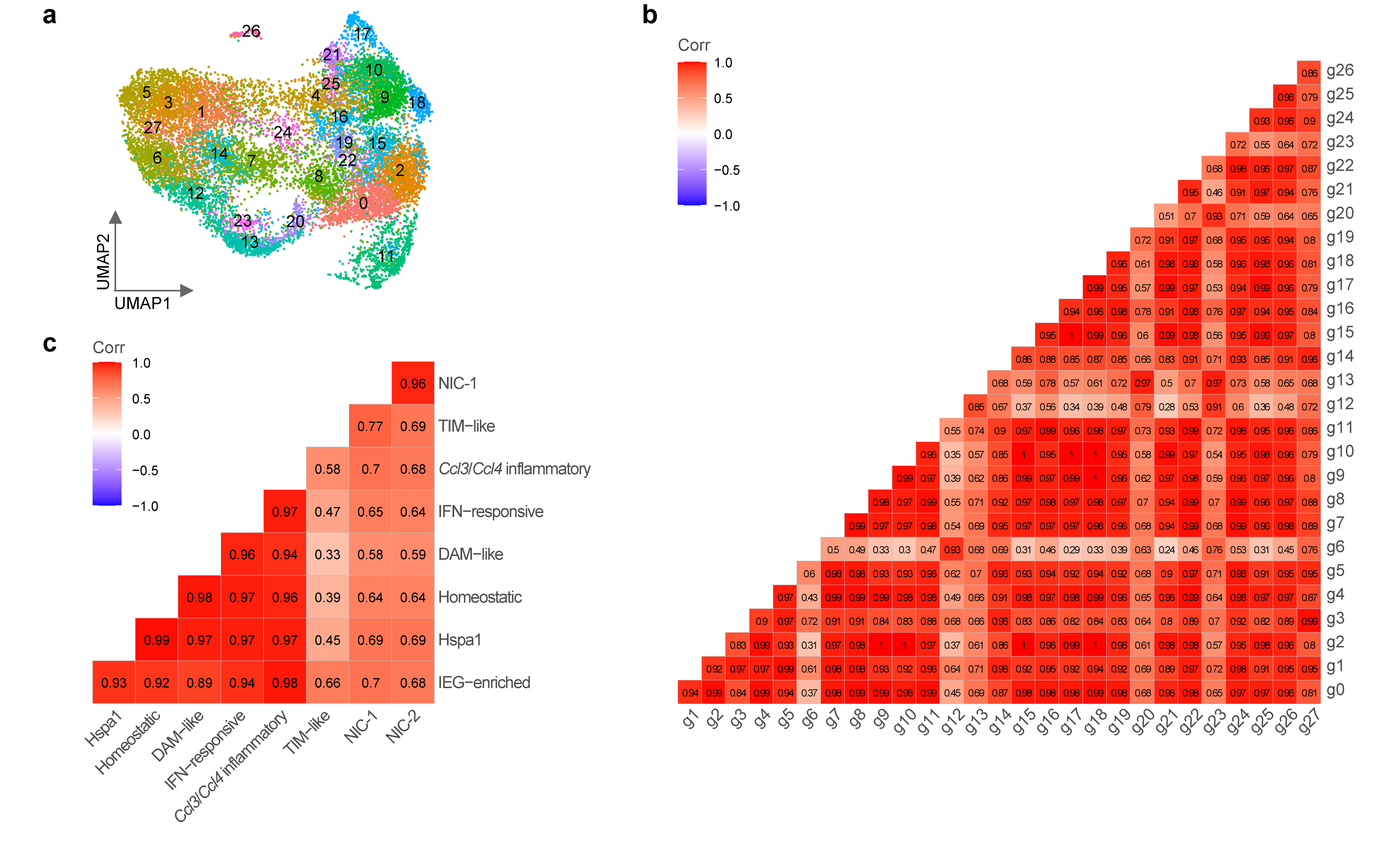

**Figure S3. Subclustering of microglia in the scRNA-seq dataset and correlations between subclusters.** **a** | UMAP plot illustrating the initial clustering result prior to subcluster merging and annotation. **b** | Pearson correlation matrix of initial microglial subclusters. **c** | Pearson correlation matrix of merged and annotated microglial subclusters.

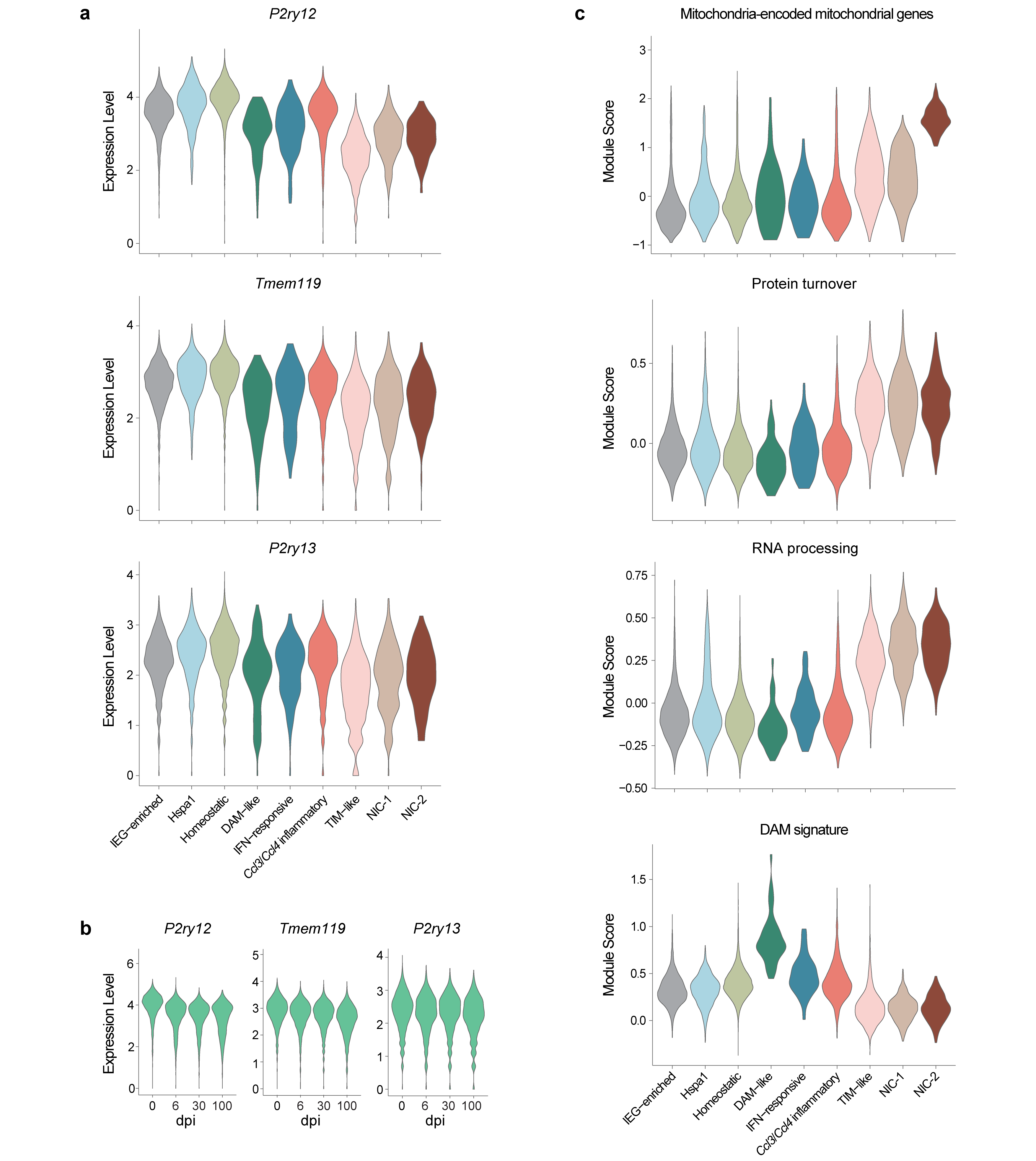

**Figure S4. Molecular signatures of microglia**. **a** | Violin plots showing the expression of microglial homeostatic genes across subclusters. **b** | Violin plots showing the expression of microglial homeostatic genes across samples. **c** | Module scores of selected gene sets across subclusters.

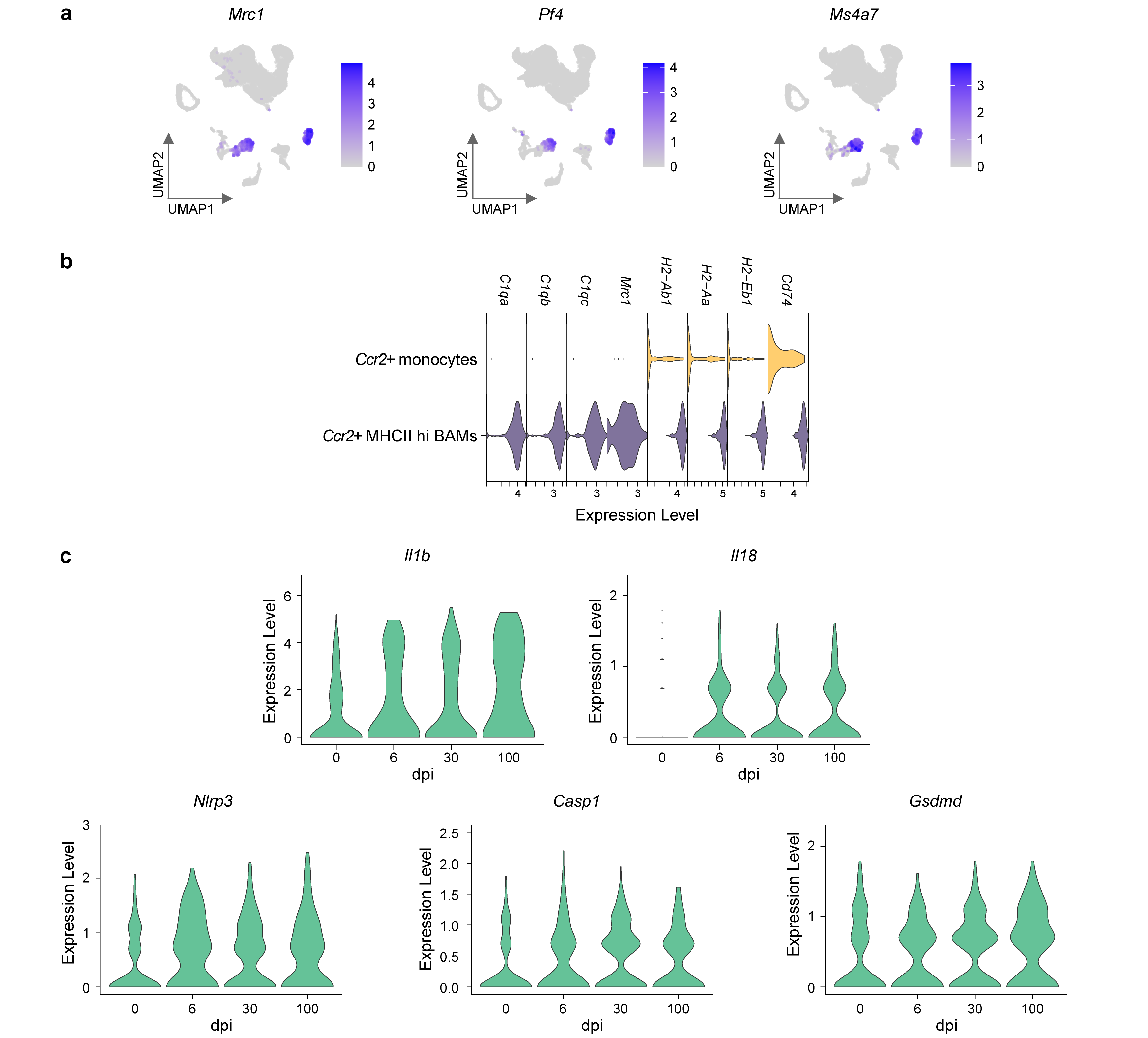

**Figure S5. Transcriptomic features of border-associated macrophages (BAMs).** **a** | FeaturePlot illustrating the expression of universal BAM signature genes. **b** | Marker genes distinguishing *Ccr2*^+^MHCII^hi^ BAMs from *Ccr2*^+^ monocytes. **c** | Expression of inflammasome-dependent cytokines and inflammasome components in MHCII^hi^ BAMs across samples.

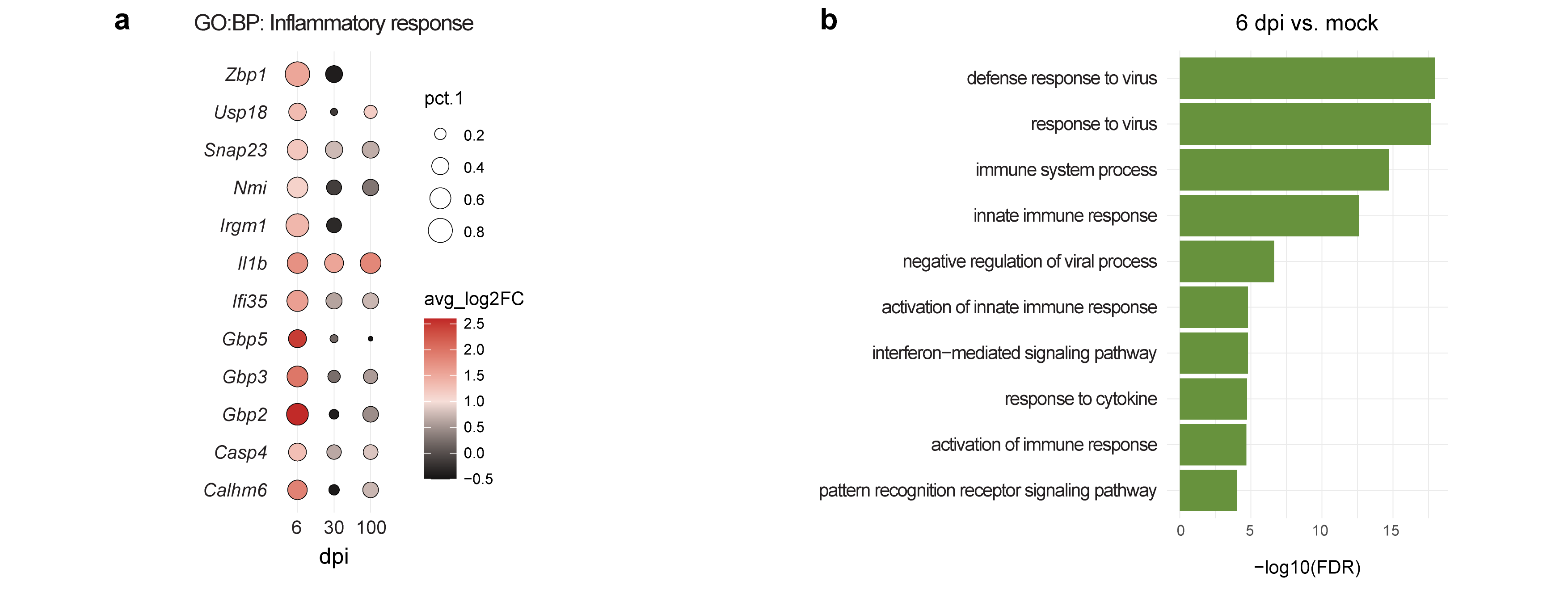

**Figure S6. Transcriptomic features of infiltrating myeloid cells**. **a** | Regulation of genes in monocytes across samples under the GO:BP term inflammatory responses. pct.1 indicates the proportion of microglia in each sample expressing indicated genes; avg_log2FC indicates the log2 fold change relative to mock. **b** | GO:BP enrichment of genes upregulated in neutrophils at 6 dpi relative to mock. Selected terms are shown.

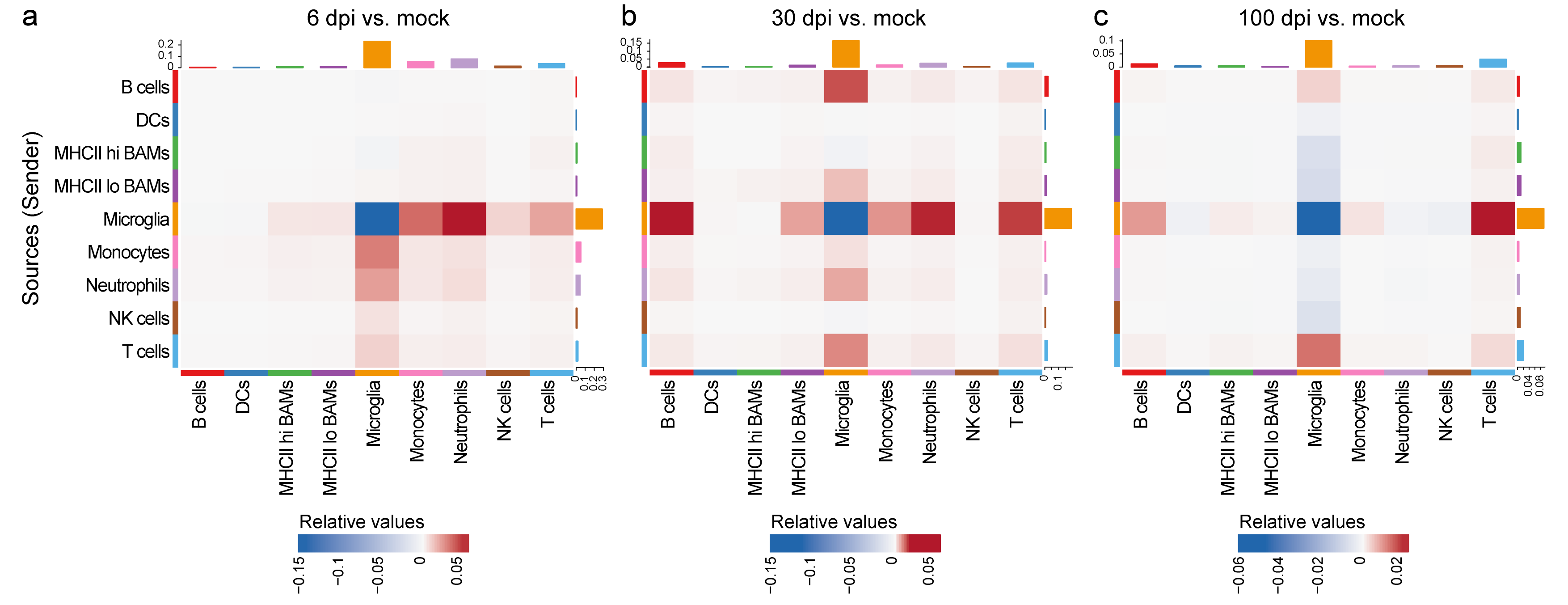

**Figure S7. Heatmaps showing cell-cell communication strength across samples, inferred by CellChat.**
